# The engulfment receptor Draper organizes the postsynaptic spectrin cytoskeleton into corrals containing synaptic proteins and promotes synaptic renewal

**DOI:** 10.1101/846121

**Authors:** Simon Wang, Mannan Wang, Hae-yoon Kim, Nicole Yoo, Matias Raski, Claire Shih, Clare Zheng, Kevin Tran, Wade Parkhouse, Charles Krieger, Nicholas Harden

## Abstract

The spectrin cytoskeleton is required for development of the Drosophila neuromuscular junction (NMJ) but its role is unclear. Here we show that the muscle spectrin lattice functions to corral membrane-associated synaptic proteins and limit their lateral mobility. Drosophila adducin, Hts, is required for integrity of the spectrin cytoskeleton and disruption of Hts function results in failure of the corrals. The spectrin cytoskeleton is itself patterned at the muscle membrane by the engulfment receptor Draper (Drpr) through regulation of Hts. We find patches of membrane where the spectrin cytoskeleton is organized into bilaterally symmetric patterns, which coincide with a field of Drpr-dependent structures similar to phagocytic pseudopods. The bilaterally symmetric patterns are likely created by folds of the muscle membrane in the pseudopods. We present evidence that the folds trap nascent boutons of motor neurons, leading to boutons with a bilaterally symmetric organization of the postsynaptic membrane. Drpr thus acts as a sensor of synaptic damage that promotes synaptogenesis.

## Introduction

The spectrin-actin cytoskeleton is required for synaptogenesis at the Drosophila neuromuscular junction (NMJ) but this role has not been extensively characterized (Blunk et al., 2014; Featherstone et al., 2001; Koch et al., 2008; Mosca et al., 2012; Pielage et al., 2008; Pielage et al., 2005, 2006). An important contributor to spectrin regulation of synaptogenesis is adducin (Hu Li Tai Shao (Hts) in Drosophila), a spectrin-binding protein that crosslinks spectrin to actin (Pielage et al., 2011; Wang et al., 2011). One route by which Hts contributes to formation of the post-synapse is through its regulation of the localization of the key scaffolding molecule Discs large (Dlg).

Here we explore further how the spectrin cytoskeleton and its regulation at the muscle membrane contribute to synapse formation. The events we consider pattern the postsynaptic membrane into three domains, the active zone, periactive zone and the subsynaptic reticulum (SSR) (Sone et al., 2000). We began by using live imaging to examine the dynamic behaviour of Dlg at the muscle membrane, which led to the finding that Dlg can move laterally along the muscle membrane, but in a regulated manner. Furthermore, Dlg adopts a distribution at the muscle membrane similar to that of α-spectrin, and as we previously demonstrated, Dlg is in close proximity to Hts as shown by the proximity ligation assay (PLA)(Wang et al., 2014). We find that other membrane-associated proteins show a similar interaction with spectrin and Hts, including mCD8-GFP, demonstrating that this interaction is sequence nonspecific. We propose that the spectrin cytoskeleton acts like a corral to restrict the movement of important synaptic proteins, thus regulating their function. A loss-of-function mutation in *hts* allows excessive interactions to occur between various synaptic proteins, likely due to destabilization of spectrin-based corrals.

In the second section of this work, we characterize the regulation of the spectrin cytoskeleton by Draper (Drpr), an engulfment receptor in phagocytosis that has been implicated in formation of the larval NMJ (Fuentes-Medel et al., 2009). We find Drpr is complexed with Hts and contributes to the formation of large gaps in the spectrin cytoskeleton throughout the muscle membrane. In patches of the muscle membrane near the NMJ, intact areas of the spectrin skeleton adopt a bilateral symmetry. Similarly, proteins at the postsynaptic membrane of the NMJ tend to show a bilateral distribution. We present evidence that the formation of Drpr-dependent pseudopods causes folding of the muscle membrane during patterning of the spectrin cytoskeleton, such that the patterns on either side of the fold are mirror images of each other. We propose that nascent boutons are trapped by the folds in the muscle membrane and mature into functional boutons with a bilaterally symmetric distribution of post-synaptic synaptic proteins. Having synapse formation dependent on pseudopod formation, provides a link between engulfment of synaptic debris and synaptogenesis, and may enable the muscle to respond to excessive synaptic damage.

## Results

### Dlg is laterally mobile in the plasma membrane of body wall muscles and this mobility is inhibited by Hts

We previously found that Hts regulates Dlg localization at the NMJ and we used FRAP to examine the effects of Hts knockdown on the dynamic behavior of this localization. We performed FRAP on Dlg-GFP in wild-type and in larvae expressing a *hts* RNAi transgene and found that reduction of *hts* significantly accelerated recovery from FRAP (Fig. 1A-D, Supplementary Movies 1 to 3). We noticed two distinct patterns of Dlg-GFP recovery in both control and Hts-knockdown body wall preparations, most commonly simultaneous restoration of Dlg-GFP in each one of a chain of bleached boutons (seen in all movies) and recovery of a bleached bouton by apparent flow from an adjacent, unbleached bouton (seen in 1 out of 8 wild-type movies, and 2 out of 10 *hts* knockdown movies)(Fig. 1A1-A3’, C1-C5’).

**Figure 1.** FRAP imaging of larval muscles expressing Dlg-GFP using mef2-Gal4 reveals that Dlg can diffuse laterally in the muscle membrane but also shows an association with membranous structures. Body wall preparations in this and subsequent figures show the muscle 6/7 pair from segment 4 of third instar larva. (A1-A3’) Imaging of NMJ of larva in which *mef2-Gal4* strain was crossed to *UAS-Dlg-GFP* stock. Note that at 3 minutes Dlg-GFP is flowing into a bleached bouton (arrowhead) from unbleached neighbor. (A2’, A3’) High magnification views are shown. Dotted line in A2’ denotes edge of bleached area. Dotted line in A3’denotes lateral flow of Dlg-GFP from unbleached bouton. See also Supplementary Movie 1. (B1-B4) NMJ of larva in which *hts* had been knocked down through expression of *htsRNAi* transgene. Dlg-GFP is restored simultaneously in the 3 bleached boutons. Note that at 3 minutes there is substantial recovery of fluorescence. See also Supplementary Movie 2. (C1-C5’) Body wall expressing *htsRNAi* in which bouton shows Dlg-GFP restoration by flow from neighboring unbleached bouton (arrowhead in C3). Panels C2’and C5’show high magnification views of flow from bouton. The dotted line in C2’marks the edge of the bleached region and in C5’ denotes the flow of Dlg-GFP into the bleached bouton. See also Supplementary Movie 3. (D) Quantification of FRAP movies showing that recovery of Dlg-GFP signal is significantly faster in *htsRNAi*-expressing larvae than in wild-type. (E) Larva in which *mef2-Gal4* strain was crossed to *UAS-Dlg-GFP* stock showing strong Dlg-GFP localization to the SSR of NMJ boutons, and nuclei. Elsewhere in the muscle membrane Dlg-GFP shows a diffuse distribution with faint puncta. (F) Enlarged view of Dlg-GFP distribution in control body wall. (G) Body wall in which Hts had been knocked down with RNAi, showing a wild-type Dlg-GFP distribution in muscle 6 on the left likely due to stochastic failure of RNAi expression. Pattern of Dlg-GFP expression in muscle 7 is consistent with association with t-tubules and overlying muscle membrane, all of which are in contact with the spectrin cytoskeleton. (H) Enlarged view of distribution of Dlg-GFP in *hts* knock down body wall preparation. Scale bars, 10 µm (A1-A4, B1-B4, C1-C6, E, G); 5 µm (A2’, A3’ C2’, C5’, F, H).

Dlg has previously been shown to be concentrated at the SSR, a structure consisting of a cluster of muscle plasma membrane folds around each bouton, and presumably the Dlg-GFP flowing into the unbleached bouton comes from this membrane store (Lahey et al., 1994). The other examples of GFP mobility we saw are also consistent with Dlg movement in the membrane and we conclude that our FRAP studies are evidence of lateral mobility of Dlg in the plasma membrane, which is accelerated when *hts* is knocked down.

### Dlg is associated with the spectrin cytoskeleton

*hts* knockdown was accompanied by a change in the appearance of extra-synaptic Dlg-GFP. Whereas in wild-type body walls, Dlg-GFP appeared as evenly scattered bright puncta, the majority of muscles where *hts* was knocked down (14/20 in ten body wall preparations) showed organization of Dlg-GFP into a network of parallel lines, many broken up (Fig. 1F-I). The punctate pattern was consistent in all samples we imaged and these results suggest an interaction of Dlg with a membrane-associated structure, which is affected by perturbation of Hts. Hts is a component of the spectrin cytoskeleton, which underlies the entire muscle plasma membrane, and we wondered if Hts was regulating Dlg distribution through its effects on spectrin. We are not aware of anyone previously characterizing the extra-synaptic spectrin cytoskeleton of the muscle membrane, probably in part due to the need to overexpose the signal at the NMJ in order to discern the extra-synaptic protein. α-spectrin and Hts staining in wild-type body walls imaged with a Zeis LSM880 with Airyscan super-resolution revealed a quite even distribution of puncta, similar to the Dlg-GFP pattern in wild-type larvae (compare Fig. 1F to Fig. 2A, B). Hts puncta showed a more detailed structure than α-spectrin, consisting of clusters of speckles. Anti-Dlg staining revealed puncta similar to α-spectrin (Fig. 2E). Strikingly, in patches of the muscle membrane near the NMJ the α-spectrin and Hts puncta showed an altered organization in which puncta appeared to have coalesced (Fig. 2C-E). This was particularly well revealed with Dlg staining, which showed a gradient of puncta morphology in the membrane (Fig. 2E). All of our stainings of the spectrin cytoskeletal puncta suggested a non-random distribution and we investigated this further using the spatial statistics 2D/3D ImageJ plugin which reveals deviation from spatial randomness in the form of clustering (attraction) or regularity (repulsion)(Andrey et al., 2010; Ollion et al., 2013). The F-function and G-function outputs from this plugin confirmed non-random distributions that were likely due to a combination of repulsion and clustering (Fig. 2F,G).

**Figure 2.**
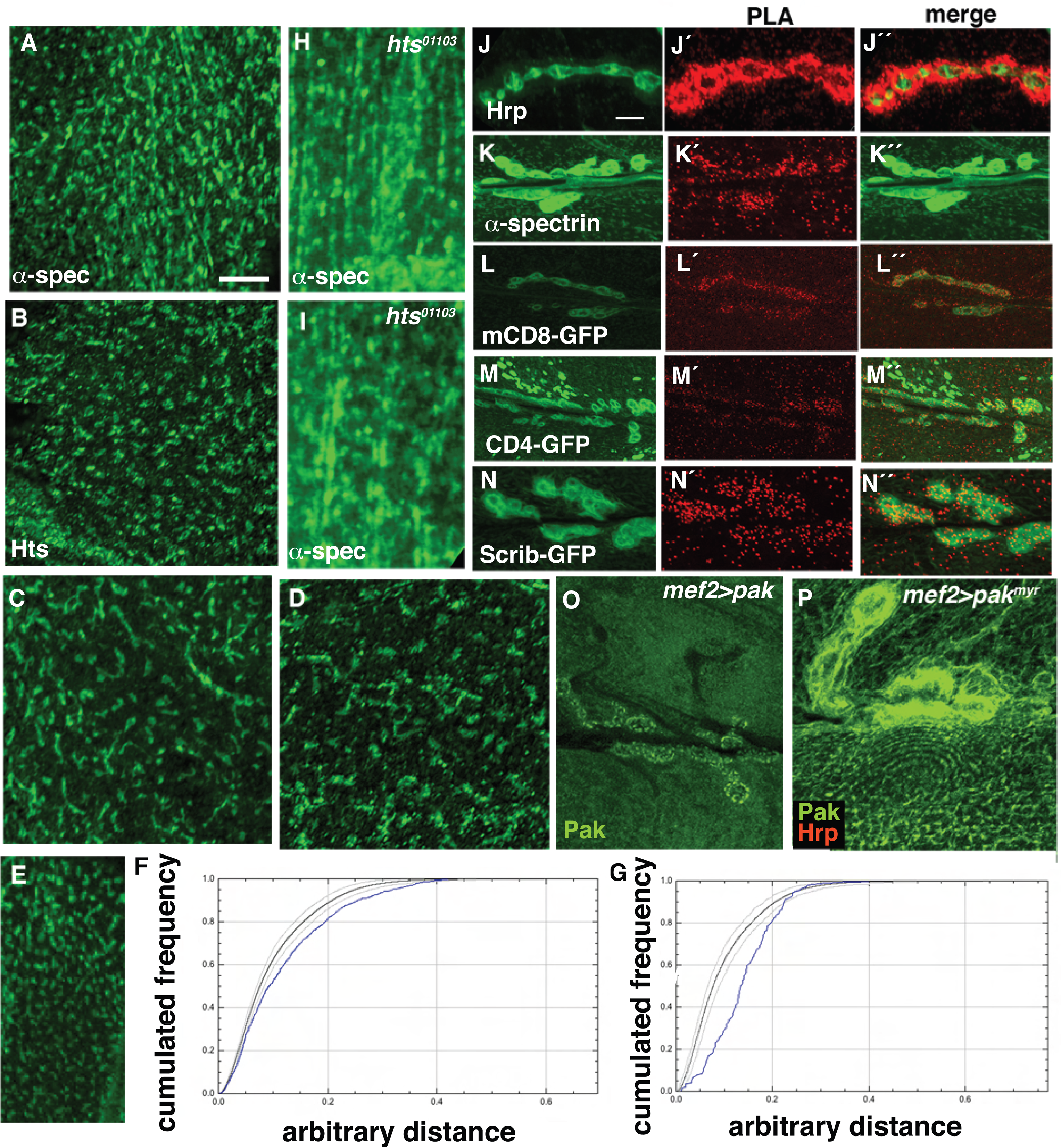
The spectrin cytoskeleton of the muscle membrane is organized into patches of high and low α-spectrin levels, and proteins associated with it show close proximity to Hts. (A, B) View of α-spectrin (A) and Hts (B) distribution in the body wall musculature of wild-type third instar larvae distant from the NMJ showing a fairly even spacing of consistently sized puncta. (C, D) View of α-spectrin (C) and Hts (D) distribution in area of muscle membrane near NMJ showing puncta of various shapes and sizes. (E) View of transition between the two types of spectrin cytoskeletal distribution shown by anti-Dlg staining. (F, G) F- and G-function plots for the puncta shown in (E). The plots, in blue, lie to the right of randomized control plots (black, 95% confidence intervals in grey) indicating non-random distribution of puncta due to both aggregation and repulsion. (H, I) Altered appearance of spectrin cytoskeleton in muscles of *hts* mutant larvae. (J-P) All panels show NMJs from third instar larvae. (J-J”) PLA signal between Hts and Dlg at SSR of each bouton. (K-K”) PLA signal between α-spectrin and Hts. (L-L”) PLA signal between mCD8-GFP and Hts. (M-M”) PLA signal between CD4-GFP and Hts. (N-N”) PLA signal between Scrib-GFP and Hts. (O) Pak over-expression in muscle using mef2-Gal4. Pak staining (green) is seen throughout the cytoplasm and at the post-synaptic active zones. (P) Expression of myristylated Pak (Pak^myr^) in muscle using mef2-Gal4. Pak^myr^ (green) accumulates in the spectrin cytoskeleton as shown by typical staining pattern in muscle membrane. It also accumulates ectopically at the SSR surrounding each bouton. Hrp staining is shown in red. Scale bars: 5 µm (A-I); 5 µm (J-P).

In *hts* mutant body walls the pattern of spectrin puncta was disrupted in favor of broken, parallel lines of spectrin staining, similar to what we saw with Dlg-GFP body wall preparations in which *hts* had been knocked down (Fig. 2H, I). Both live imaging and immunohistochemistry suggested that Dlg was associating with the spectrin cytoskeleton throughout the muscle membrane, and this association with spectrin could provide an additional means for Hts regulation of Dlg.

In the well-characterized red blood cell (RBC), the spectrin cytoskeleton forms a lattice under the plasma membrane (Matsuoka et al., 2000). It has been suggested that such a spectrin-lattice is present at the post-synapse where it is involved in synaptic organization (Pielage et al., 2006). Studies in the RBC have demonstrated that the spectrin cytoskeleton forms corrals that restrict the movement of the protein Band 3 and it has been proposed that corralling could be a general mechanism for regulating the distribution of membrane proteins (Tomishige et al., 1998). We suspected that Dlg is corralled in the spectrin cytoskeleton, where it will come into close proximity with Hts, as we previously demonstrated with PLA (Wang et al., 2014) and Fig. 2J-J”. We found that Hts is in close proximity to spectrin in body walls, as expected (Fig. 2K-K”). There is no sequence-specific requirement for corralling in the body wall spectrin cytoskeleton, as we found that expression of two versions of membrane-localized GFP accumulated in a spectrin-like pattern and showed a PLA signal with Hts (Fig. 2L-M”). One of these reporters, mCD8-GFP, has previously been reported to label both the SSR and T-tubules, modifications of the plasma membrane, and presumably is distributed throughout the entire muscle plasma membrane when expressed in muscle (Fujita et al., 2017). In contrast, cytoplasmic GFP did not localize to either the SSR or T-tubules, suggesting that membrane association is required for corralling. Dlg is in a membrane-associated protein complex with the Scrib protein at the NMJ, and we found that Scrib-GFP, like Dlg, is embedded in the spectrin cytoskeleton where it is in close proximity to Hts as demonstrated by a PLA signal (Mathew et al., 2002)(Fig. 2N-N”).

Taken together, our live imaging results, combined with our analysis of the spectrin cytoskeleton, suggest that a patterned distribution of spectrin corrals at the muscle membrane immobilize membrane-associated proteins including the key synaptic scaffolding molecule Dlg.

### Pak can be redirected to spectrin corrals and the SSR by addition of a myristylation tag

We have shown that exogenous GFP can be directed to the spectrin cytoskeleton by targeting it to the plasma membrane, but can endogenous NMJ proteins be similarly localized by membrane targeting? The serine/threonine kinase Pak is an NMJ protein but did not show a spectrin-like distribution in the muscle membrane, indicating that it is not normally accumulating in the spectrin corrals (Fig. 2O). In the absence of an upstream signal Pak occurs as inactive dimers in the cytoplasm but can be constitutively localized to the membrane by addition of a myristylation tag (Pak^myr^) (Hing et al., 1999; Parrini et al., 2002).

We overexpressed wild-type Pak in the muscle using the mef2-Gal4 driver and saw diffuse, cytoplasmic localization as well as typical Pak enrichment at the postsynaptic side of the active zones (Fig. 2O). Strikingly, Pak^myr^ expressed using mef2-Gal4 showed a distribution similar to that of Dlg, accumulating in suspected spectrin corrals as shown by the spectrin-like Pak staining pattern throughout the muscle membrane, and high levels of Pak in the SSR (Fig. 2P). We conclude from this experiment, and the above results with GFP reporters, that membrane association is a key factor in proteins being corralled by the spectrin cytoskeleton.

### Dlg released from spectrin corrals shows increased interactions with NMJ proteins

We predicted that disruption of the spectrin corrals would liberate Dlg for increased interaction with NMJ proteins. Dlg is localized in both the SSR and the periactive zone of NMJ synapses and shows functional interactions with a number of active zone proteins, including stabilizing glutamate receptors containing the subunit GluRIIB (Chen and Featherstone, 2005; Sone et al., 2000). Conversely, the kinase Pak is required for Dlg accumulation at the NMJ (Parnas et al., 2001). Using PLA, we found that Dlg was in close proximity to both GluRIIB and Pak, presumably at the periactive zone-active zone boundary, indeed much of the PLA signal with Pak was outside the Hrp-stained region of the boutons (Fig. 3A-A”, C-C’). In *hts* mutant body walls, where the spectrin cytoskeleton is compromised, there was an increase in PLA signals between Dlg and GluRIIB and Pak, indicating that more of these proteins were available for interactions (Fig. 3B-B”, Fig.3 D-D”, Fig. S1A, B). In a search for other Dlg-interacting proteins we looked for an interaction between Dlg and Drpr, an important regulator of synaptic growth that co-localizes postsynaptically with Dlg and found that there was a strong, periactive zone localization of PLA signals between the two proteins, which spread throughout the muscle membrane in *hts* mutants (Fuentes-Medel et al., 2009)(Fig. 3E-F”, Fig. S1C). If this phenotype is due to disruption of spectrin corrals, it should also be seen with knockdown of α-spectrin and we saw widespread PLA between Dlg and Drpr following RNAi knockdown of α-spectrin in the muscle (Fig. 3G-G”, Fig. S1D).

**Figure 3.**
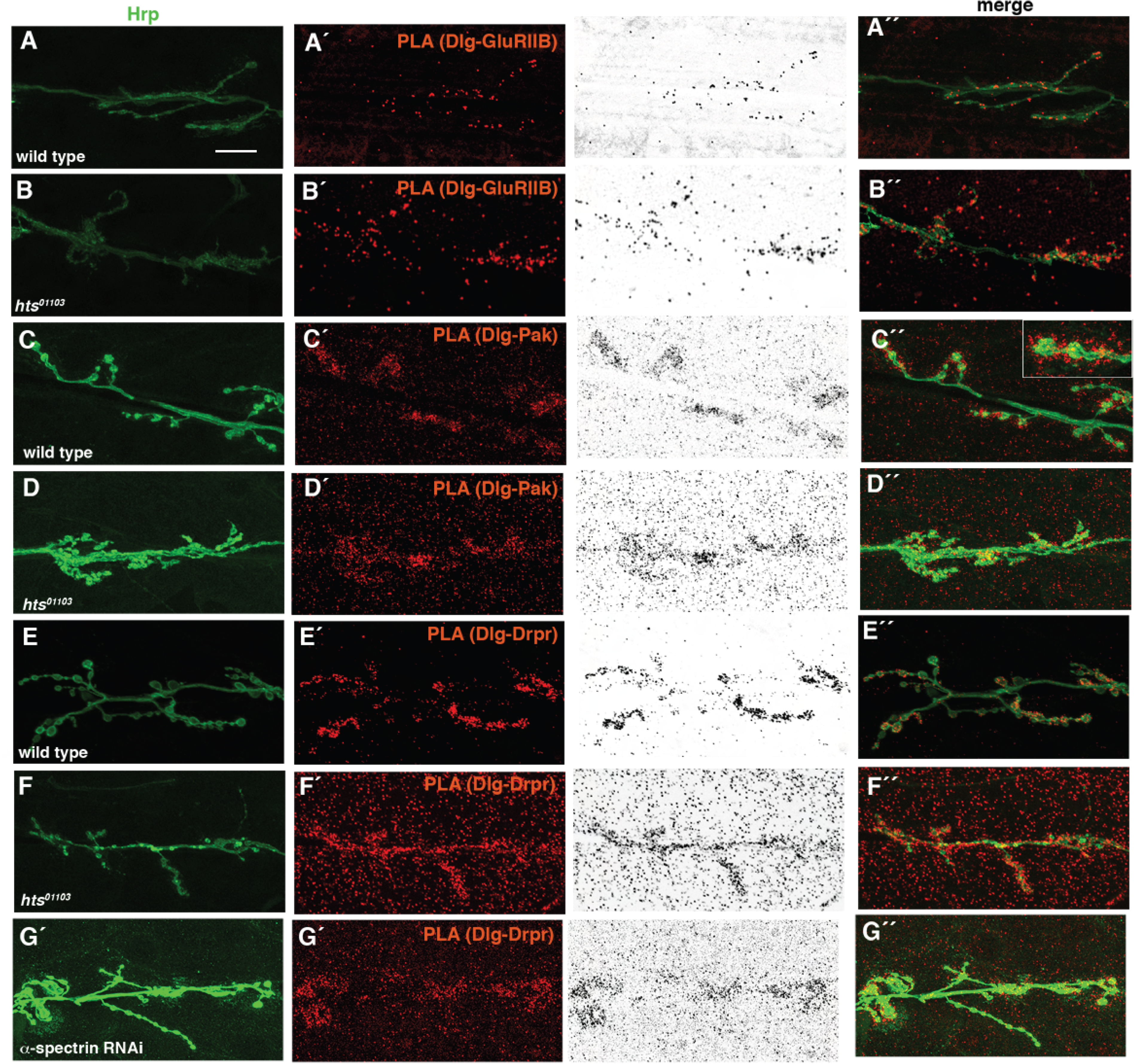
Dlg shows excessive interactions with other NMJ proteins in *hts* mutant larvae. All panels show NMJs and surrounding muscle membrane from third instar larvae. Inverted images of PLA are included for better visualization of signal. (A-B”) PLA between Dlg and GluRIIB in wild type (A-A”) and *hts* mutant (B-B”) larvae. (C-D”) PLA between Dlg and Pak in wild type (C-C”) and *hts* mutant (D-D”) larvae. Inset in panel CC’ shows PLA signal around the boundary between periactive and active zones. (E-G”) PLA between Dlg and Drpr in wild type (E-E”), *hts* mutant (F-F”) and α-spectrin knockdown larvae (G-G”). Scale bar, 20 µm.

Taken together, our live imaging and PLA studies indicate that availability of Dlg for interaction with other proteins is regulated by its confinement to spectrin corrals.

### Evidence that Drpr inhibits Hts in patterning of the spectrin cytoskeleton

As noted above, Dlg liberated from spectrin corrals interacts with the synaptic protein Drpr, and we wondered if Drpr might be involved in Dlg regulation. In addition to looking for direct interactions between Drpr and Dlg (unpublished results) we looked for an interaction between Drpr and Hts, an established regulator of Dlg, using PLA and found that the two proteins formed complexes scattered around the muscle membrane, with the PLA signals often occurring in doublets and triangular triplets (Fig. 4A, B). This pattern is distinct from the other Hts-associated proteins we have examined, which have a distribution that closely matches that of α-spectrin. We compared the distribution of Drpr-Hts PLA signals and Drpr immunohistochemistry with that of Dlg and α-spectrin in the muscle membrane and found that Drpr was consistently inside or at the edges of the areas of reduced α-spectrin and Dlg staining (Fig. 4A-B”). This Drpr distribution appeared to be non-random and we assessed this using the 2D/3D ImageJ plugin, which confirmed that the pattern was a non-random distribution consistent with repulsion (Fig. S2).

**Figure 4.**
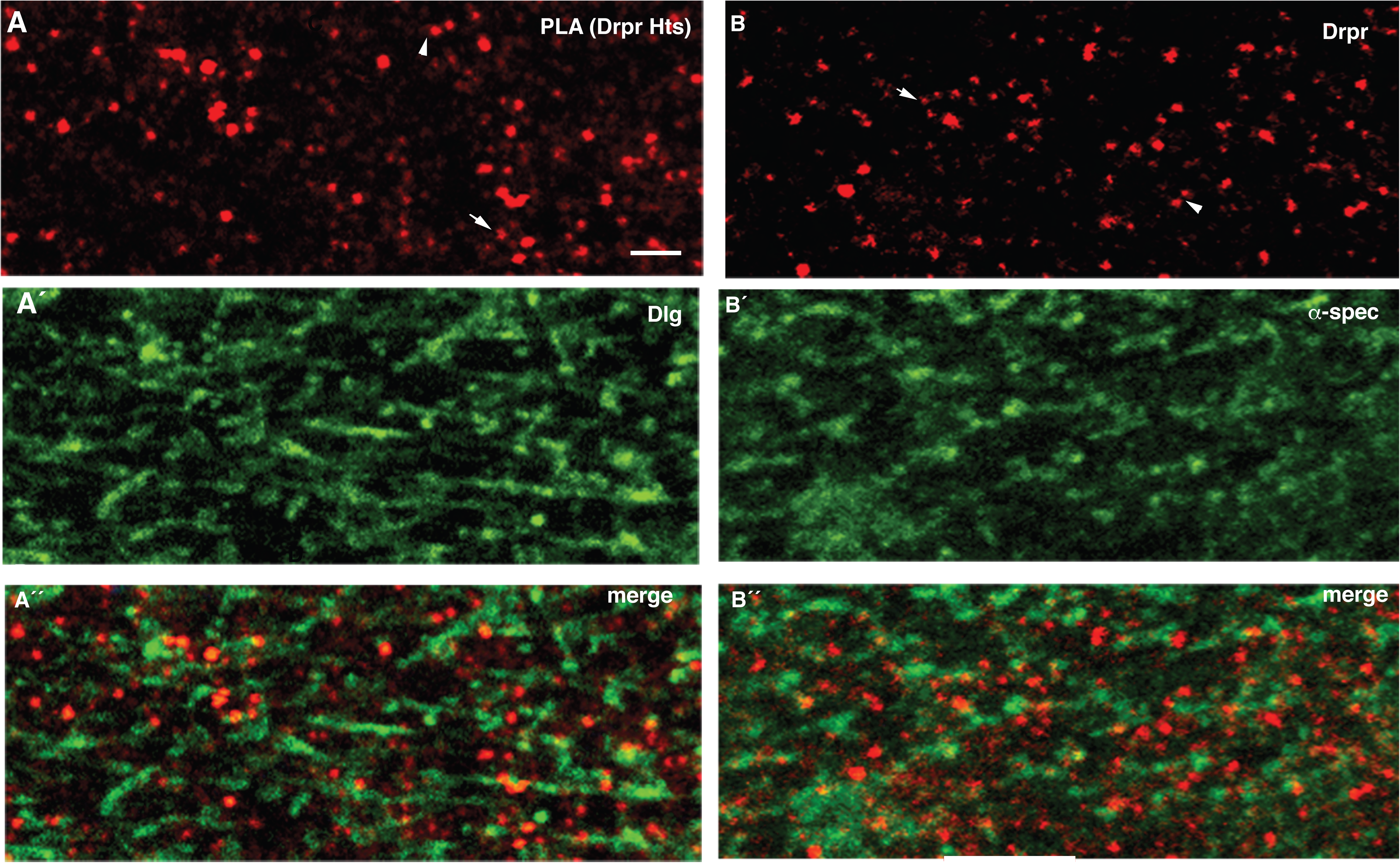
The muscle membrane spectrin cytoskeleton has Drpr-associated gaps. All panels show body walls of wild-type third instar larvae. (A, B) PLA between Hts and Drpr (A) or staining for Drpr (B) show tendency to form doublet signals (arrowheads) and triangular triplet signals (arrows). (Á-B”) Co-staining in green for the spectrin cytoskeleton markers Dlg (A’, A”) and α-spectrin (B’, B”) reveals that Drpr localizes in or at the edge of gaps in the spectrin cytoskeleton. See Fig. S2 showing that F- and G-function plots of Drpr puncta demonstrate a non-random distribution consistent with repulsion. Scale bar, 1 µm.

### Parts of the muscle membrane with a bilateral symmetry of protein distribution may be favored as post-synaptic membranes for synaptogenesis

As noted above, in parts of the muscle membrane near the NMJ the spectrin cytoskeleton showed a different organization from elsewhere in the extra-synaptic muscle membrane (Fig. 2C-E). Close examination revealed organization of the cytoskeleton in these regions into roughly bilaterally symmetric assemblies, with each assembly being around 4 to 9 µm across and each one being distinct (Fig. 5, Fig. S3). No assemblies of this size were seen when images of puncta were reorganized using the Pattern Maker Plugin in Adobe Photoshop, indicating that these assemblies are unlikely to be occurring by chance (Fig. S3D, F). In the intact spectrin cytoskeleton, Hts directly interacts with α-spectrin and, as can be seen in Fig. 5A, B there were PLA signals between Hts and spectrin in the bilaterally symmetric patches of cytoskeleton but not in the gaps, even though PLA is very sensitive. This result confirms that the gaps are indeed regions of little or no spectrin cytoskeleton.

**Figure 5.**
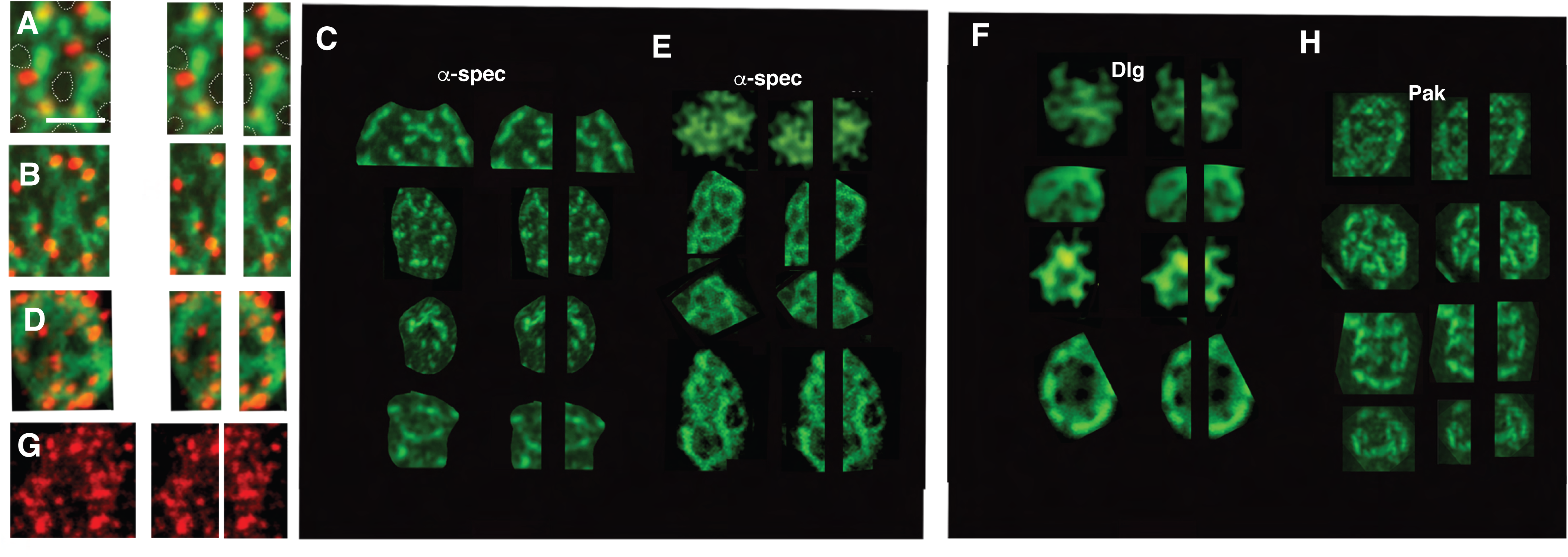
Both the muscle plasma membrane and the post-synaptic membrane of boutons can show a bilaterally symmetric organization of synaptic proteins. (A, B) Views of the spectrin cytoskeleton in the muscle plasma membrane showing bilateral symmetry. α-spectrin is shown in green, PLA for Hts and α-spectrin is shown in red. Gaps in the cytoskeleton are outlined in A. To the right of each panel is a view split along the axis of symmetry to facilitate evaluation of degree of symmetry. (C) Examples of bilaterally symmetric assemblies seen in muscle plasma membrane. See Fig. S3 for examples of such assemblies in situ. (D) Post-synaptic membrane of bouton, stained as in (A, B). (E, F) Further examples of bouton post-synaptic membranes stained with anti-α-spec or anti-Dlg. (G) Post-synaptic membrane of bouton stained with anti-Drpr. (H) Post-synaptic membrane of bouton stained with anti-Pak. Note that patterns are more precise than with anti-Dlg or anti-α-spectrin. Scale bar, 2 µm (A, B, D, G).

The gaps in Dlg and α-spectrin staining seen throughout the muscle membrane cytoskeleton were similar to the gaps in Dlg and α-spectrin seen in the postsynaptic membrane where boutons of motor neurons synapse with the muscle to form the NMJ (Lahey et al., 1994; Pielage et al., 2006)(Fig. 5D-F). Similar to the muscle membrane, the postsynaptic membrane synapsing with many type 1B boutons showed a similarly sized (5 µm across) bilateral symmetry in PLA between Hts and α−spectrin, and staining for α-spectrin, Dlg and Drpr (Fig. 5D-G). In general, the bilateral patterns were simpler and more imperfect and distorted at the post-synaptic membrane, with a noticeable exception being Pak, which showed quite complex bilateral patterning (Fig. 5H). It was not possible to quantify this phenotype because many boutons could not be evaluated due to their orientation in the body wall preparation, but bilaterally symmetric synapses were readily detectable. These results suggest that the muscle membrane is pre-patterned for function as the postsynaptic membrane before contact is made with boutons.

To characterize further Drpr interaction with the spectrin cytoskeleton of the muscle membrane we looked at two *drpr* alleles, the previously characterized null allele *drpr^Δ5^* and *drpr^MB06916^*, which is a transposon insertion in an intron of *drpr* (Freeman et al., 2003). We began by staining body wall preparations with anti-α-spectrin and looking for effects on patterning and density of α-spectrin staining. *drpr^MB06916^* and *drpr^Δ5^* mutants exhibited an altered pattern of spectrin staining, shifting to a significantly denser distribution with no bilateral symmetry (Fig. 6A-F, Fig.S4). The elevated spectrin seen in *drpr* mutants persisted when these alleles were placed over a deficiency, *Df(3L)BSC181*, removing the *drpr* locus, indicating that these were *drpr* loss-of-function phenotypes (Fig. S5). Given our model of corralling of proteins such as Dlg by the spectrin cytoskeleton, we looked at Dlg distribution in *drpr* mutants and observed an increased density of Dlg at the muscle membrane for the *drpr^MB06916^* allele but not the *drpr^Δ5^* allele (Fig. 6D-F, Fig. S4). We next examined the distribution of Pak, which localizes specifically to the post-synaptic active zone (Sone et al., 2000). Both *drpr* alleles had NMJs with normally localized Pak, but 100% of *drpr^MB06916^* mutants (n=20) exhibited scattered ectopic accumulations of Pak throughout the body wall (Fig. 6G, H, I-I”). We noticed that ectopic Pak accumulated adjacent to high levels of Dlg expression, and that this phenotype was seen when either *drpr^MB06916^* or *Df(3L)BSC181* were placed over a wild-type chromosome indicating that it was a dominant loss-of-function phenotype (Fig. S5). When *Df(3L)BSC181* was placed over the *drpr^Δ5^* allele chromosome the ectopic Pak phenotype disappeared, suggesting dominant suppression by the *drpr^Δ5^* genetic background (Fig. S5B).

**Figure 6.**
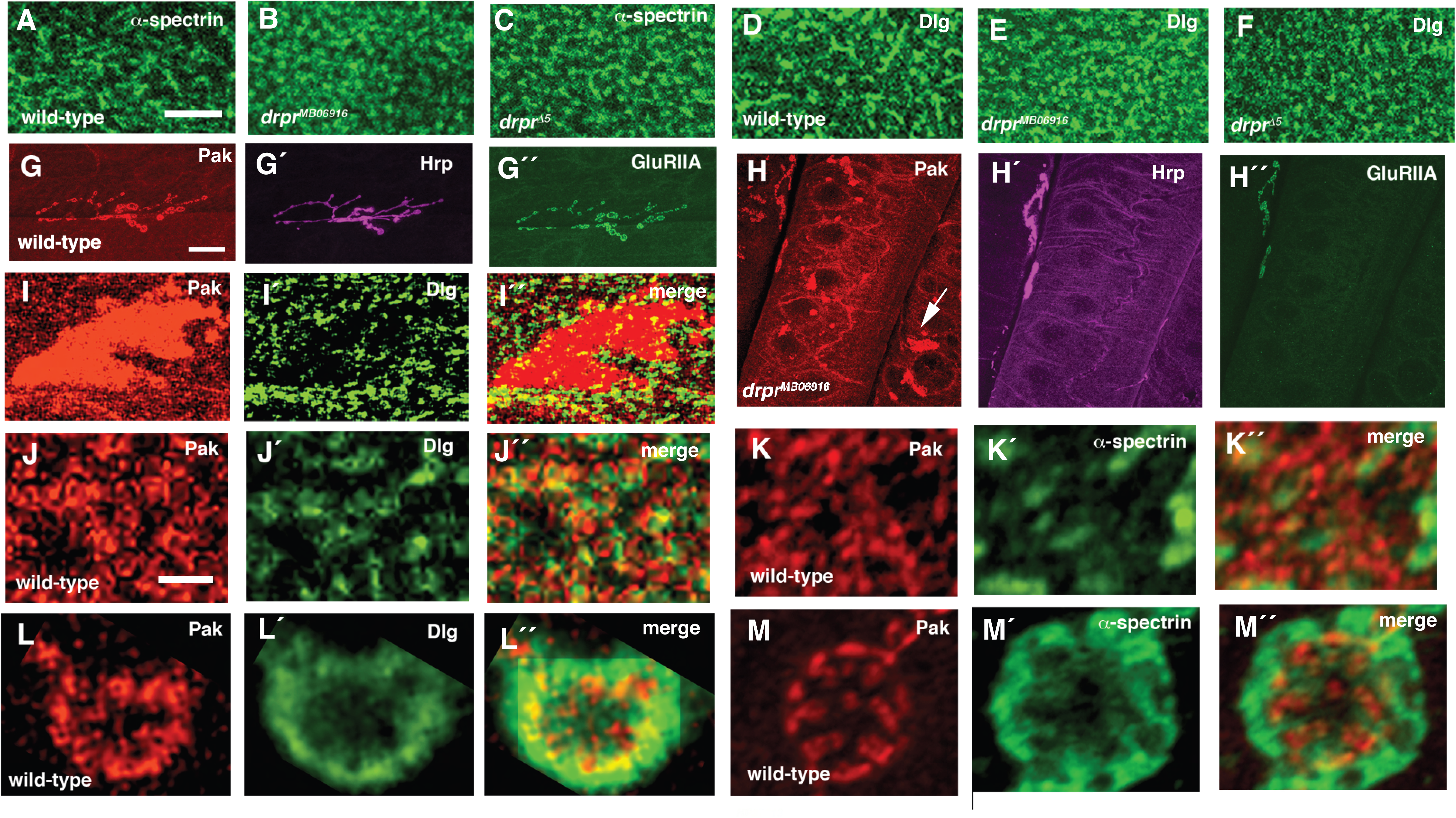
Perturbation of Drpr function has effects on the spectrin cytoskeleton and distribution of synaptic proteins. (A-C) α-spectrin distribution in wild-type (A) and *drpr* mutant body walls (B, C). The spectrin cytoskeleton is denser in *drpr^MB06916^* and *drpr^Δ5^* mutants. (D-F) Dlg distribution in wild-type (D) and *drpr* mutant (E, F) body walls. Dlg is denser than wild-type in *drpr^MB06916^* but not *drpr^Δ5^* mutants. (G-G”) Wild-type NMJ stained for Pak (G), Hrp (G’) and GluRIIA (G”). (H-H”) *drpr^MB06916^* mutant body wall, stained as in G-G”, showing scattered, ectopic accumulations of Pak. (H) A large accumulation of Pak is marked by an arrow. Staining for Hrp (H’) and GluRIIA (H”) reveal an NMJ at top left of the panel, but demonstrate that ectopic accumulations of Pak are not in NMJs. (I-I”) High magnification view of ectopic patch of Pak (I) in *drpr^MB06916^* mutant co-stained for Dlg (I’); the merge (II”) demonstrates that Pak tends to accumulate where Dlg levels are low. (J-M”) Wild-type body walls. Pak (J) does not show a lot of colocalization with Dlg (J’) or the spectrin cytoskeleton (K’) either in the extra-synaptic muscle membrane (J-K”) or at the bouton (L-M”). Note the similarities in protein distributions between the extra-synaptic muscle membrane and the bouton. Scale bars: 10 µm (A-F, I-I”), 15 µm (G-H”), 1 µm (J-M”).

We looked at the relationship between Pak and Dlg in wild-type larvae and found that, consistent with the Dlg-Pak PLA signal (Fig. 3Ć), the Pak distribution overlapped partially with that of Dlg (Fig. 5J-J”, Pearson’s correlation coefficient 0.37). We suspected that the overlapping distributions of Pak and Dlg were in areas where Dlg had been released from spectrin corrals, and consistent with this Pak showed little overlap with areas of strong spectrin staining (Fig. 5K-K”, Pearson’s correlation coefficient 0.12). Again, we observed similarities between the extrasynaptic muscle membrane and the postsynaptic side of boutons, with Pak at the postsynaptic membrane showing some overlap with Dlg but little with spectrin (Pearson’s Correlation Coefficients 0.44 and 0.24 respectively) (Fig. 5L-M”).

To summarize, the patterning of the center of the postsynaptic membrane, inside the ring of the SSR, was similar in appearance to the extra-synaptic muscle membrane.

### Drpr-dependent pseudopods may drive the formation of bilateral symmetry in the spectrin cytoskeleton

The most remarkable observation we made with regard to the spectrin cytoskeleton was the presence of patches of bilateral symmetry. We considered membrane folding as a possible cause as it could bring regulatory complexes into contact with the spectrin cytoskeleton on the opposing membrane of the fold. Drpr is a member of a family of engulfment receptors that phagocytoses synaptic debris and immature boutons as part of the normal development of the NMJ (Fuentes-Medel et al., 2009). Work on the C. elegans Drpr ortholog CED-1 indicates that this phagocytosis involves the formation of phagocytic cups with pseudopod extensions (Shen et al., 2013). We examined the surface of wild-type body walls stained for Dlg or α-spectrin and found that 95% (55/58) of wild-type individuals exhibited what appeared to be a series of intersecting extensions of the muscle membrane which contained pseudopod-like structures with an unusual morphology of “prongs” at the extracellular end (Fig. 7A1-B2). We tested various antibodies to see if they could label the membrane extensions and found that they were strongly stained by an antibody against Pretaporter (Prtp), a ligand for Drpr that we have yet to characterize for a possible function in the muscle membrane (Fig. 7C)(Kuraishi et al., 2009). Based on the various stainings and viewing the structures from different perspectives, we concluded that they were fields of shallow phagocytic cups, sitting above the muscle contractile apparatus (Fig. 7A1-E). Fuentes-Medel et al previously showed that muscle cells take up synaptic debris and degrading boutons resulting from synaptic plasticity, and we determined if the cups we discovered were used in this uptake (Fuentes-Medel et al., 2009). Indeed, these structures were found to encapsulate spherical bodies that were stained with anti-Hrp and antibodies against GluRIIB, both bouton markers (Fig. 7D-G). The phagocytic cups in C. elegans are CED-1-dependent structures, and to determine if the phagocytic cups in Drosophila were similarly dependent on Drpr, we examined the effects of gains or losses of Drpr function on these structures. In larvae over-expressing full length Drpr in muscle, the cups were significantly elongated (p=0.0029), being on average twice as long in the plasma membrane-cytoplasm axis as in the plane of the epithelium (n=41), compared to wild-type in which the ratio was 1.5 (n=46)(Fig. 7H-H’). The phagocytic cups were rarely or not seen in *drpr^MB06916^* or *drpr^Δ5^* mutants or larvae expressing a *drprRNAi* transgene in muscle, with a frequency of 0.8 % (2/26), 0% (0/34), and 0% (0/23), respectively (Fig. 7I, J).

**Figure 7.**
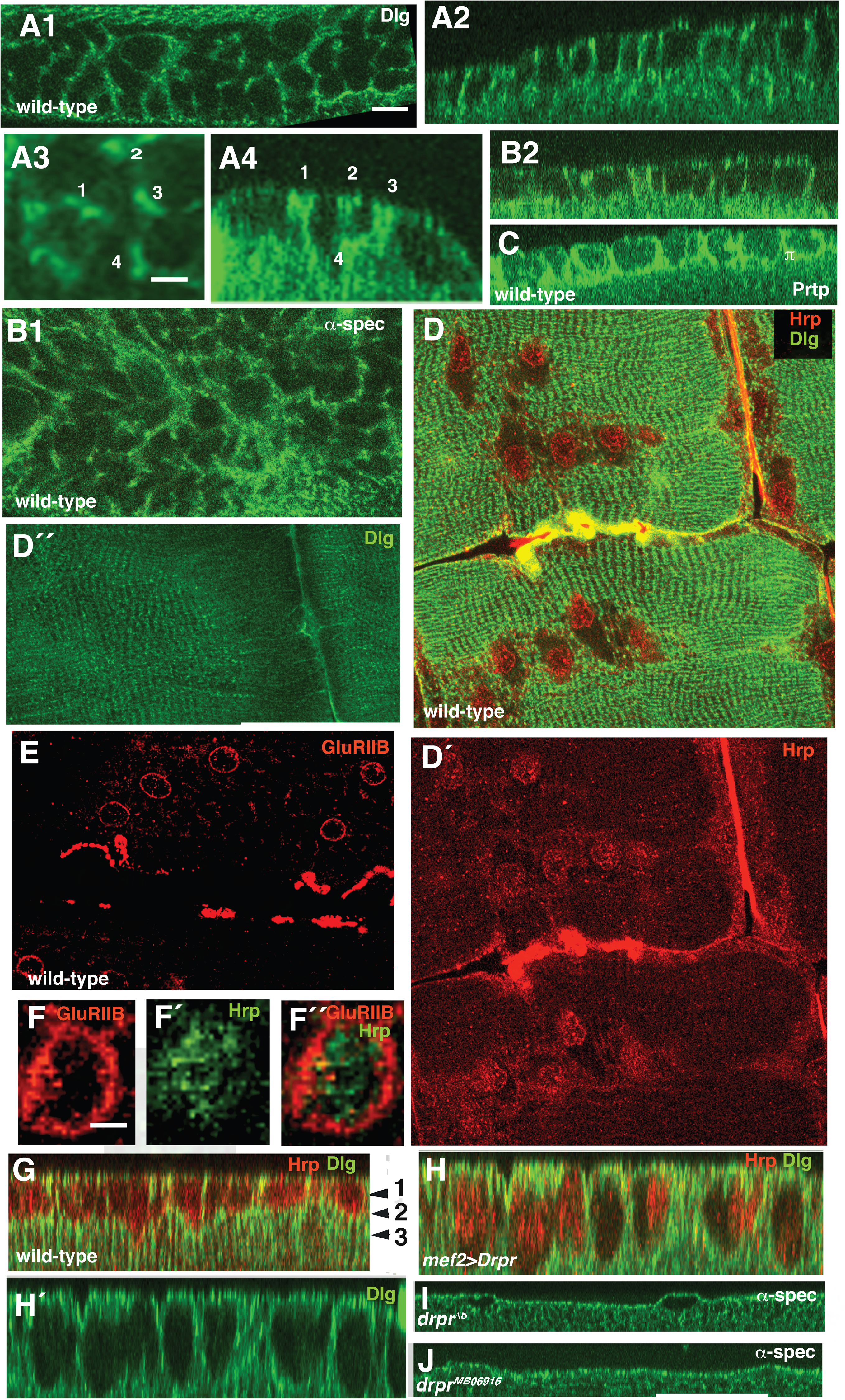
Pseudopod-like extensions of the muscle plasma membrane may form capture sites for nascent boutons through invaginations created by membrane folding. (A1, A2) En face (A1) and optical cross-sectional (A2) views of membrane ridges in in a wild-type body wall stained for Dlg. Pseudopod-like structures protrude from muscle membrane, with each bright spot on the membrane ridge in A1 being one pseudopod (A3, A4). Higher magnification views of same body wall preparation focused on one cluster of 4 pseudopods. (B1, B2) En face (B1) and optical cross-sectional (B2) views of membrane extensions in a wild-type body wall stained for α-spectrin, revealing the same structures as Dlg staining. (C) Muscle plasma membrane staining with anti-Prtp showing the pseudopods to be wave like structures. (D-D”) Wild-type body wall preparation stained for Dlg (green) and Hrp (red) showing indentations of the muscle plasma membrane containing Hrp-positive material. (D’) shows that indentations contain spherical objects that are likely degenerating boutons. D” shows intact contractile apparatus. (E-E”) Degenerating bouton positive for Hrp (green) and GluRIIB (red). (G) Cross-sectional view generated from Z stack indicating that indentations are cup-like structures containing Hrp-positive material. Arrowheads 1, 2 and 3 denote the different focal planes used in this figure, corresponding to panels A1, A3, and B1; D, D’ to F; and D”, respectively. (H, H’) Optical cross-sectional view of body wall in which Drpr had been over-expressed in muscle showing elongated phagocytic cups. (I, J) Optical cross-sectional views of *drpr* mutant body walls stained for α-spectrin. There are no pseudopod extensions. Scale bars: 10µm (A1, A2, B1-D’, E), 5µm (A3, A4) 2µm (F-F”).

### Perturbation of Drpr function disrupts the formation of ghost boutons

We suspected that the phagocytic cups were involved in creation of the bilaterally symmetric assemblies, and consistent with this, assemblies were detected in flat areas of membrane amongst the pseudopods (Fig. 8A, C). A feature of normal body wall preparations are neurites bearing ghost boutons, which are boutons lacking post-synaptic markers such as Dlg (reviewed in (Ataman et al., 2008; Fuentes-Medel et al., 2009; Menon et al., 2013). We define a ghost neurite as a thin, Hrp-positive, Dlg-negative process from which ghost boutons form. We found ghost neurites extending across regions of bilateral symmetry in wild-type body wall preparations and noticed that the smaller boutons sat adjacent to the bilaterally symmetric assemblies (Fig. 8B), whereas rare, larger ghost boutons were centered over an assembly (Fig. 8D-D”). Through Z-sectioning, we determined that the ghost boutons te<u>nded to be adjacent to</u> phagocytic cups with 76 % (25/33) of ghost boutons in 8 individuals so positioned (Fig. 8B-F”). Given this, we wondered if their formation might be dependent on Drpr-mediated phagocytosis and we looked for an effect of perturbing Drpr function on ghost boutons. Typically, we saw between 0 and 2 ghost neurites for each muscle 6/7 pair, regardless of genotype. In wild type these ghost neurites all had ghost boutons, whereas ony 28% and 20% of *drpr^Δ5^* or *drpr^MB06916^* ghost neurites, respectively, had ghost boutons (Fig. 8G, H). Similarly, individuals expressing a *drprRNAi* transgene in muscle had no ghost boutons (Fig.8I). We over-expressed Drpr in muscle and again saw a loss of ghost boutons, suggesting that either gains or losses of Drpr could disrupt ghost bouton formation (Fig. 8J).

**Figure 8.**
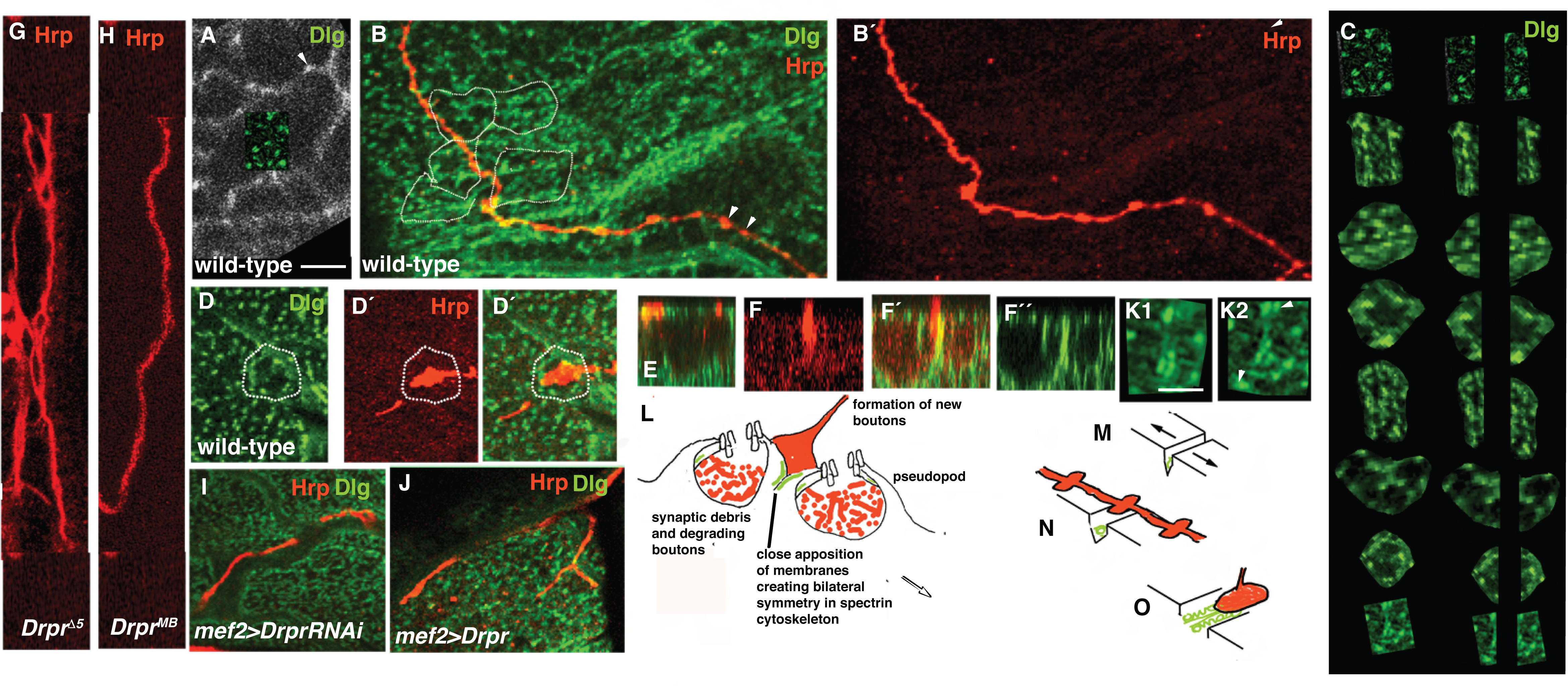
Ghost boutons are associated with regions of bilateral asymmetry and are dependent on Drpr for their existence. Unless otherwise stated all samples were stained with anti-Hrp (red) and anti-Dlg (green). (A) Example of bilaterally symmetric distribution of Dlg in the vicinity of pseudopods in a wild-type body wall. Gray scale image shows membrane ridges supporting pseudopods (bright spots, arrowhead). Image in green is a different focal plane showing that bilaterally symmetric Dlg assembly is located above the membrane ridges. This assembly is also shown in panel C. (B) Ghost bouton neurite adjacent to bilaterally symmetric assemblies. Dashed line outlines some of the assemblies, which are also shown in panel C. (B’) Neurite shown on its own, showing ghost boutons as swellings. (C) Examples of bilaterally symmetric assemblies seen in this figure. To the right of each image is a view split along the axis of symmetry to facilitate evaluation of degree of symmetry. (D-D”) Wild-type ghost bouton centered over a bilaterally symmetric assembly. (E-F”) Z-sections showing ghost boutons adjacent to phagocytic cups. Boutons are solid red masses whereas synaptic debris in cups is particulate. (F) is a cross-sectional view of the two ghost boutons marked with arrowheads in (B). (G-J) Losses or gains of Drpr function perturb ghost bouton formation. *Drpr* mutants (G, H) or individuals expressing an RNAi transgene against Drpr or larvae over-expressing Drpr in muscle all show reduced numbers of ghost boutons on ghost neurites. (K1,K2) Live imaging of bilaterally symmetric spectrin cytoskeleton as revealed by mCD8-GFP expression. Panels are images taken 15 minutes apart showing emergence of bilaterally symmetric pattern. Arrowheads mark the emergence of puncta. See also Movie 4.s(L-O) Model of proposed mechanism of Drpr-dependent formation of new boutons in response to phagocytosis. (L) Emergence of Psuedopods forming phagocytic cups leads to fold in plasma membrane. As cups fill with synaptic debris neighboring pseudopods move in opposite directions to close up membrane. (M) Pulling by pseudopods (arrows) opens up fold in membrane to form a cleft. (N) Cleft captures nascent ghost boutons. (O) Bouton matures as membrane unfolds to reveal bilateral symmetry of spectrin cytoskeleton and synaptic proteins, forming post-synaptic membrane of bouton. Scale bars: 5µm (A-B’, D-D’, G-J), 3µm (K, K’).

The results on ghost boutons suggested that their presence was dependent on regions of the membrane where pseudopods are creating phagocytic cups. Especially striking was the tendency of small ghost boutons to be inserted in the membrane between phagocytic cups or between bilaterally symmetric assemblies, whereas larger ghost boutons, and mature boutons in the NMJ, tend to be centered over bilaterally symmetric assemblies.

Staining of the pseudopods with antibodies labelling the spectrin cytoskeleton presumably only reveals their internal structure and they were wave-shaped as revealed by Prtp staining of the plasma membrane (Fig.7C). We measured their width at the base to see if they could likely accommodate a folded region big enough for a symmetric pattern of around 4 to 9 µm^2^ across split between the two sides of the fold. With an average width of 8.26 µm (n=10), the pseudopods would appear to be large enough to supply the folded membranes that develop a bilateral symmetry of the spectrin cytoskeleton. We observed the dynamic behavior of the bilaterally symmetric assemblies by live imaging body walls expressing mCD8-GFP and found that a complete pattern emerged over time, for example one of a pair of mirror image dots would be visible before the other (Fig. 8C, K1-K2, Movie 4).

In summary, Drpr may have two roles in organizing the spectrin cytoskeleton. First, it promotes gaps in the spectrin cytoskeleton throughout the muscle membrane and second, membrane folding between Drpr-dependent pseudopods results in a bilaterally symmetric organization of the gaps in the cytoskeleton.

## Discussion

### A corralling model for spectrin regulation of NMJ development

Previous studies have demonstrated a requirement for the spectrin-actin cytoskeleton in proper postsynaptic development at the larval NMJ; our work provides mechanistic insight into this requirement (Blunk et al., 2014; Featherstone et al., 2001; Koch et al., 2008; Mosca et al., 2012; Pielage et al., 2008; Pielage et al., 2005, 2006). We have demonstrated that the spectrin lattice can corral membrane-associated proteins such as Dlg and segregate them from ectopic interactions with other NMJ proteins. Consistent with this, a previous study demonstrated scattering of Dlg in β- or α-spectrin mutants (Featherstone et al., 2001). In live imaging studies, we have visualized the lateral movement of Dlg along the muscle membrane, as well as a pattern of static Dlg localization consistent with an association with the spectrin cytoskeleton throughout the muscle membrane including the SSR. The pattern of Dlg localization into broken parallel lines is consistent with an association with the cytoskeleton of the t-tubules, extensions of the muscle plasma membrane that Dlg had previously been shown to localize to (Safi et al., 2016). We propose that loss of *Drosophila* adducin, Hts, results in mobilization of Dlg from the corrals and this phenotype may mimic a normal regulatory mechanism in which localized inactivation of Hts is used to “open” the corrals and release their contents. This is supported by our FRAP result showing that Dlg can flow out of the SSR even in wild-type body walls. A recent study of the role of the spectrin cytoskeleton in Drosophila myoblast fusion demonstrates another function for spectrin corralling in muscle development: the corralling of the cellular adhesion molecules Dumbfounded and Antisocial in embryonic myoblasts (Duan et al., 2018). In wild-type embryos Dumbfounded and Antisocial are tightly clustered but become dispersed in α/β_H_-spectrin double mutants. Similarly, E-cadherin distribution in vertebrates appears to be regulated by corralling by the spectrin cytoskeleton and depletion of adducin in bronchial epithelial cells results in increased mobility of E-cadherin-GFP (Sako et al., 1998) (Abdi and Bennett, 2008).

Previous studies have considered the sources of Dlg at the dynamic NMJ (Thomas et al., 2000; Zhang et al., 2007). Thomas et al. describe the movement of Dlg during development from a subcortical network to the postsynaptic membrane and this requires domains in the Dlg protein (Thomas et al., 2000). For example, deletion of the HOOK domain disrupts Dlg localization at both the subcortical network and at the synapse. Our results indicate that this subcortical network is the spectrin cytoskeleton and suggest that HOOK deletion prevents Dlg retention in the corrals due to insufficient membrane localization. An earlier FRAP study suggested that Dlg-GFP fluorescence was not replenished from the SSR of neighboring boutons, but our own FRAP data indicate that Dlg can move laterally along the plasma membrane including between adjacent boutons (Zhang et al., 2007).

### Evidence that Drpr, through inhibition of Hts, contributes to areas of disrupted spectrin cytoskeleton

Drpr likely negatively regulates Hts in Drpr-Hts complexes. Disruption of a subset of Hts molecules could lead to localized collapses of the spectrin cytoskeleton and the resulting large gaps. The spacing of Drpr and the spectrin islands is consistent with a non-random distribution caused by repulsion. Interestingly, a mouse homolog of Drpr, MEGF10, is required for the orderly arrangement of retinal neurons through a process of homotypic repulsion (Kay et al., 2012). Cells expressing MEGF10 repulse each other, leading to an orderly, non-random arrangement. At the muscle membrane the distribution of Hts complexes could be dictated by both repulsion from Drpr and by the spacing of Hts molecules on the spectrin lattice and this will lead to evenly-distributed gaps in the spectrin cytoskeleton. Our analysis suggested aggregation in addition to repulsion in organization of the spectrin cytoskeleton, and the folds in the plasma membrane that we propose could affect the overall distribution of the spectrin cytoskeleton through protein clustering. Huge gaps in the spectrin cytoskeleton have been described at the lateral membrane of MDCK cells, where spectrin is confined to motile microdomains covering about half the membrane (Jenkins et al., 2015). Tropomodulin1 is a support protein that, like adducin, caps actin filaments and in so doing stabilizes the spectrin cytoskeleton. The RBC spectrin cytoskeleton in *tropomodulin 1* mutant mice is full of large gaps and is similar in appearance to the areas of disrupted muscle membrane cytoskeleton of Drosophila larvae, supporting the idea that Drpr is affecting a support protein or proteins (Moyer et al., 2010).

### A model for guidance of post-synaptic membrane formation by the spectrin cytoskeleton

We propose a model for the participation of the spectrin cytoskeleton in formation of the postsynaptic membrane at the Drosophila NMJ. Drpr complexes with Hts, leading to a “sea” of disrupted cytoskeleton at the muscle membrane with “islands” of intact spectrin cyoskeleton. These islands corral various membrane associated proteins such as Dlg and contain the nascent zones of the post-synaptic membrane of any bouton that contacts them. The post-synaptic membrane consists of the SSR, the periactive zone, and the active zone with the latter two organized in the muscle membrane prior to the arrival of the bouton. Pak, which is a major regulator of postsynaptic active site formation, is attracted by high levels of Dlg in the island corrals, probably contacting Dlg through the spectrin fence, but also Dlg released from the corrals by transient breaks in the cytoskeleton (Albin and Davis, 2004; Lee and Schwarz, 2016; Parnas et al., 2001; Ramos et al., 2015; Wang et al., 2016). This interpretation is based on our PLA and colocalization studies indicating close proximity between Pak and Dlg, and the finding that excessive amounts of Pak are recruited to the muscle membrane adjacent to high levels of Dlg in *drpr^MB06916^* mutants and larvae bearing the *Df(3L)BSC181* deficiency. The interaction between Pak and Dlg at the NMJ is bidirectional in that Dlg localization at the NMJ requires Pak (Parnas et al., 2001). Why do only *drpr^MB06916^* mutants show ectopic accumulations of Pak? The likely reason for this not occurring with the *drpr^Δ^*^5^ allele is that it doesn’t cause a significant increase in Dlg levels in the muscle membrane.

Although *drpr* mutants have reduced numbers of boutons they survive larval development, and although their spectrin cytoskeleton is denser, it still has gaps which may be sufficient for synaptic function. There might be redundancy between Drpr and other regulators of the spectrin cytoskeleton, furthermore, as discussed below, Drpr may be largely concerned with synaptic renewal rather than with the initial formation of NMJs.

### Folds in the muscle membrane caused by formation of Drpr-dependent pseudopods may function as sites for capture of nascent boutons

Although the gaps in the spectrin cytoskeleton of the Drosophila muscle membrane are similar to those in other systems, the bilateral symmetries we see in parts of the muscle membrane of the fly are to our knowledge unique. These symmetries occur in regions rich in Drpr-dependent pseudopodia, and we propose that the symmetries are contained in folds produced by neighboring pseudopods growing out from the plasma membrane (see model in Fig. 8L-O). The bilaterally symmetric pattern within these folds might result from a homophilic cell-cell adhesion molecule producing the same distribution on both sides of the fold. If the adhesion molecule stabilizes or blocks the spectrin cytoskeleton, the pattern of the spectrin cytoskeleton will be dictated by that of the adhesion molecule and will be mirrored in the membrane on the other side of the fold. For the mirror images to be visible, the fold must open up, which could reveal the image in stages. Consistent with an unfolding, our live imaging showed that an entire bilaterally symmetric pattern is not visible all at once. We propose that as the pseudopods of neighboring phagocytic cups extend in opposite directions during phagocytosis, a cleft opens between them which is derived from a fold. At some point in the opening of the cleft it may be large enough to trap a neurite with a nascent ghost bouton. Ghost boutons have previously been described as activity dependent immature boutons that can develop into mature structures (Ataman et al., 2008). Our work suggests that in addition to activity Drpr dependent phagocytosis contributes to ghost bouton formation; the evidence that these do become mature boutons being the persistent bilateral organization of proteins in many boutons at the NMJ. It is not unexpected that an invagination should be able to trap a neurite, as there are numerous examples of synapses in which the pre-synaptic terminal invaginates into the post-synaptic process, including the Drosophila NMJ (Petralia et al., 2017; Prokop, 1999). Indeed, during initial NMJ development in the Drosophila embryo, boutons are embedded in a shallow indentation in the muscle (Prokop, 1999). Phagocytosis has been proposed as a mchanism for invagination of a pre-synaptic terminal into the post-synaptic process; our work supports an indirect role for phagocytosis (Petralia et al., 2017).

In *drpr* mutants, there are still spectrin islands but they are no longer organized in a bilaterally symmetric pattern. The islands of spectrin cytoskeleton at the muscle membrane may also provide the foundation of the SSR, as they are enriched in SSR proteins. Past1 is a protein involved in development of the SSR and has been described as localizing to a“postsynaptic tubular-reticular domain”, which we believe is the spectrin cytoskeleton (Koles et al., 2015).Thus, Past1 may be corralled similarly to Dlg and can readily be delivered for SSR development.

We have proposed mechanisms by which the spectrin cytoskeleton underlying the muscle membrane can guide the formation of an orderly array of active zones on boutons at the NMJ. The bilaterally symmetric patterning of the spectrin cytoskeleton is more precise at the extra-synaptic muscle membrane than at the post-synaptic membrane of the bouton. This is almost certainly due to presence of the SSR at the bouton, and it is noteworthy that Pak, a protein that doesn’t normally localize to the SSR and is not corralled, can show quite a precise bilaterally symmetric distribution at the post-synaptic membrane of the bouton. The bilaterally symmetric spectrin cytoskeleton might not have any functional relevance, simply being a side effect of fold formation, or it may have some bearing on synaptic function. One possibility is that it could be used to achieve balanced adhesion of the bouton to the muscle membrane and in support of this a recent study demonstrated that the spectrin cytoskeleton acts as a nanoscale ruler mediating cell-cell adhesion between cells derived from neural stem cells (Hauser et al., 2018). It will be of interest to determine how widely corralling is used in creating patterned membranes; given the ubiquitous nature of the spectrin cytoskeleton this could be occurring in many cell types.

Our work opens up exciting new directions of research on the spectrin cytoskeleton and joins a body of literature demonstrating that the spectrin cytoskeleton is far from simply a passive cellular structure but rather an active participant in developmental and cellular events. Through our characterization of Drpr, a regulator of the spectrin cytoskeleton, we have discovered a potential feedback mechanism in mature larvae that ensures that loss of neurons in development is not excessive. The more engulfment occurring the greater the neurite trapping and de novo bouton formation (Fig. 8L). Furthermore, new boutons form where they are required as it is engulfment of old boutons that creates the capture sites for the new boutons. Thus, Drpr acts as a sensor of synaptic damage that can promote synaptic renewal.

## Materials and Methods

### Fly Stocks

All stocks and crosses were incubated at 25°C. *w^1118^* was used as a wild-type control strain. *UAS-eGFP-dlgA* was a gift from U. Thomas (Koh et al., 1999), *drpr^Δ5^*, *UAS-drprRNAi* and *UAS-drprI* from M. Freeman (Freeman et al., 2003), and *UAS-scrib-GFP* from D. Bilder (Zeitler et al., 2004). RNAi transgenic lines against Hts and α-spectrin were from the Vienna *Drosophila* Resource Center, and were tested for knockdown of target genes as shown in Fig. S6. All other stocks were from the Bloomington *Drosophila* Stock Center.

### Larval Body Wall Preparation

Body wall dissections of crawling third instar larvae were performed as previously described (Wang et al., 2015). Mutant stocks were re-balanced over GFP-containing balancers to distinguish non-fluorescent, homozygous mutant larvae from their fluorescent, heterozygous siblings. For transgenic analysis, homozygous *UAS*-transgene-bearing males were crossed to homozygous *Gal4*-bearing virgin females ensuring that all progeny carried one copy of each. Immunohistochemistry was performed as previously described (Ramachandran and Budnik, 2010; Wang et al., 2011), with primary antibodies used described in the next section.

### Proximity Ligation Assay

PLA was performed as previously described (Wang et al., 2015). Dissected body walls were blocked with 1% BSA for one hour in pre-siliconized microcentrifuge tubes. In a given experiment, all genotypes, which were distinguished by unique cuts made to the corner of each body wall, were incubated within a single tube, thus receiving identical processing. The body walls were then incubated overnight at 4°C with rabbit and mouse primary antibodies against the two proteins of interest in 1% BSA, along with a 1:100 dilution of goat anti-Hrp antibody (Jackson ImmunoResearch – 805-605-180) to mark neuronal membranes. The following primary antibodies were used for PLA and/or immunohistochemistry: 1:50 mouse anti-Brp (Developmental Studies Hybridoma Bank – nc82) (Wagh et al., 2006)1:100 mouse anti-Dlg (DSHB – 4F3) (Parnas et al., 2001), 1:500 rabbit anti-Drpr (gift from M. Freeman) (Freeman et al., 2003), 1:200 rabbit anti-GFP (Sigma-Aldrich – G1544), 1:50 mouse anti-GluRIIA (DSHB – 8B4D2) (Kurusu et al., 2012)1:250 rabbit anti-GluRIIB (gift from A. DiAntonio) (Marrus et al., 2004), 1:100 rabbit anti-Hrp (Sigma-Aldrich - G1544), 1:10 mouse anti-Hts (DSHB – 1B1) (Zaccai and Lipshitz, 1996), 1:200 rabbit anti-HtsM (gift from L. Cooley)(Petrella et al., 2007), 1:200 rabbit anti-Pak (Harden et al., 1996), 1:50 rat anti-Prtp (Gift from Y. Nakanishis) (Kuraishi et al., 2009 and 1:10 mouse anti-α-Spec (DSHB – 3A9) (Dubreuil et al., 1987). After three washes with PBT for ten minutes each, the body walls were incubated with a 1:5 dilution of anti-rabbit PLUS (Sigma-Aldrich – DUO92002) and anti-mouse MINUS (Sigma-Aldrich – DUO92004) PLA probes in 1% BSA for 90 minutes at 37°C. Samples were then washed twice with Wash A for five minutes each, and incubated in Ligation solution (Sigma-Aldrich – DUO92008) for one hour at 37°C. Following two washes with Wash A for two minutes each, the body walls were incubated in Amplification solution (Sigma-Aldrich – DUO92008) for two hours at 37°C. The body walls were then washed twice with ten minute washes of Wash B followed by a single wash with 0.01× Wash B for one minute. To detect the goat anti-Hrp antibody, the body walls were subsequently incubated with a 1:200 dilution of a FITC-conjugated anti-goat secondary antibody (Jackson ImmunoResearch – 705-095-003) in 1% BSA for one hour. The body walls were then washed three times for ten minutes each in PBT and stored in VECTASHIELD (Vector Laboratories – H-1200) at −20°C until ready for imaging. For negative controls, experiments were done with either one primary antibody omitted or done in larvae mutant for one the two proteins (Fig. S6). Images of muscle 6/7 NMJs from abdominal segment 4 were taken on either a Nikon A1R laser scanning confocal microscope with NIS-Elements software or a Zeiss LSM800, with experimental samples and their controls imaged under identical acquisition settings. All images were processed with Adobe Photoshop.

### Fluorescence Recovery After Photobleaching

Third instar larval body walls were dissected and mounted onto slides using HL3 saline solution (Stewart et al., 1994). Photobleaching of synaptic boutons innervating muscles 6/7 on abdominal segment four was performed on a Quorum Wave FX Spinning Disc Confocal System using Volocity software. Stacks were taken every 10 seconds (i.e. six time points per minute) post bleaching. All imaging was completed within 30-60 minutes of dissection of live animals.

### Image analysis

In order to evaluate protein distributions in the extra-synaptic muscle membrane, equivalent brightness adjustments were made to all images from a given experiment using the white input slider of the levels adjustment function of Photoshop. As this brightness adjustment generally caused overexposure of NMJ protein staining, care was taken to only analyze extra-synaptic muscle membrane signals not immediately adjacent to the NMJ.

To quantify the density of extrasynaptic muscle membrane protein deposition, brightened images were opened in ImageJ, inverted and converted to binary. The measure function was then used to determine the percent area of the image that had black pixels.

We used the 2D/3D Spatial Analysis plugin to evaluate the distribution of Drpr puncta in the extra-synaptic muscle membrane. Three possible patterns for puncta are evaluated with this plugin: completely random, aggregated due to attraction, or regular (non-random) due to repulsion. Confocal images of Drpr puncta were opened in ImageJ, inverted and converted to binary. The plugin was then used to calculate the G-function and the F-function plots. The G-function is the cumulative distribution function of the punctum to closest punctum distance, whereas the F-function is the cumulative distribution function of the arbitrary point to closest punctum. The plugin also calculates the average G- and F-function plots for randomized, control versions of the Drpr puncta. A significant deviation from non-randomness is visualized as a plot lying outside the 95% confidence interval for the randomized control plot.

For evaluating colocalizations, we used the Coloc2 plugin of ImageJ to calculate Pearson’s correlation coefficients.

## Acknowledgments

We thank D. Bilder, M. Freeman and U. Thomas for fly stocks; L. Cooley, A. DiAntonio, M. Freeman and Y.Nakanishis for antibodies and T. Audas for comments on the manuscript. This work was funded by grants from the Canadian Institutes of Health Research, the Natural Sciences and Engineering Research Council of Canada (NSERC) and the Rare Diseases: Models & Mechanisms Network to NH and NSERC and ALS Canada/Brain Canada to CK.

