## Supplementary Data for "The engulfment receptor Draper organizes the postsynaptic spectrin cytoskeleton into corrals containing synaptic proteins and promotes synaptic renewal"

### Supplementary Figure Legends

#### Movies

Movie 1. Dlg-GFP-expressing body wall. Movie is shown at 200X faster than actual time.

Movie 2. Body wall expressing Dlg-GFP and *htsRNAi*. Movie is shown at 20X faster than actual time.

Movie 3. Body wall expressing Dlg-GFP and *htsRNAi*. Movie is shown at 20X faster than actual time.

Movie 4. Body wall expressing mCD8-GFP. Movie is shown at 10X faster than actual time.

**Figure S1.** Quantification of PLA data comparing Dlg interactions with other NMJ proteins in wild-type larvae versus *hts* mutant larvae (A-C) or wild-type larvae versus $\alpha$-spectrin knockdown larvae (D). n numbers are shown on the graphs. *, $p<0.05$; **, $p<0.010$; ***, $p<0.001$.

**Figure S2**. (A) Anti-Drpr staining of body wall preparation used to evaluate Drpr distribution using spatial statistics 2D/3D ImageJ plugin.(B) F-function showing Drpr plot (blue) to left of randomized control plot (black, 95% confidence intervals in grey). G-function showing Drpr plot (blue) to right of randomized control plot (black, 95% confidence intervals in grey. Both plots indicate distribution by repulsion.

**Figure S3.** Examples of bilaterally symmetric patches in an $\alpha$-spectrin-stained muscle membrane from wild-type larvae. Dotted lines outline such patches. Panels D and F are scrambled images of panels C and E, respectively. Scale bar, 5µm.

**Figure S4.** Quantification of effects of *drpr* alleles on $\alpha$-spectrin and Dlg. n numbers are shown on the graphs. *, $p<0.05$; **, $p<0.010$; ***, $p<0.001$.

**Figure S5.** All body wall preparations stained with anti-Pak (red). (A-C) *drpr* spectrin cytoskeleton phenotypes of increased density and loss of bilateral symmetry (insets show $\alpha$-spectrin stain) are confirmed when alleles are placed over a deficiency, *Df(3L)BSC181*, removing the *drpr* locus. (D) Similar to *drpr*$^{MB06916}$ heterozygotes, the ectopic Pak phenotype is seen in *Df(3L)BSC181* heterozygotes but is suppressed in *Df(3L)BSC181/drpr*$^{\Delta 5}$ individuals (B).

**Figure S6.** Controls for PLA and RNAi experiments. (A-B′) Hts PLA experiments done in a *hts* mutant background generate no signals. (C, C′) A Dlg PLA experiment done in a *dlg* mutant background generates no signals. (D) Control larva in which *mef2Gal4* driver line had been outcrossed to *w*$^{1118}$ wild-type strain, showing robust Hts (green) staining in SSR and extrasynaptic muscle membrane. (E) Expression of HtsRNAi in the muscle eradicates postsynaptic Hts leaving neuronal, presynaptic Hts (green) that colocalizes with Hrp (magenta). (F) Control larva in which *mef2Gal4* driver line had been outcrossed to *w*$^{1118}$ wild-type strain, showing robust $\alpha$-spectrin (green) staining in SSR and extra-synaptic muscle membrane. (G) Expression of $\alpha$-spectrin RNAi in the muscle eradicates postsynaptic Hts leaving neuronal, presynaptic Hts (green) that colocalizes with Hrp.

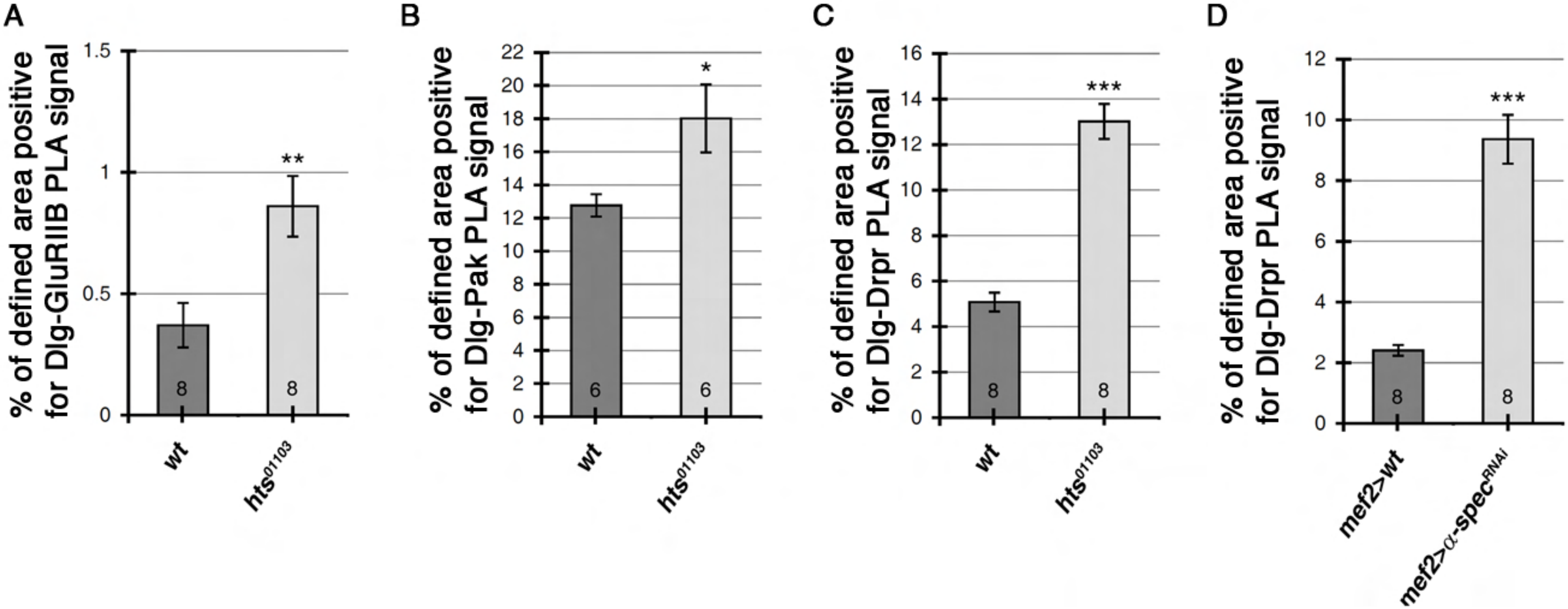
A
% of defined area positive for Dlg-GluRIIB PLA signal
1.5
1
0.5
0
**
8
8
wt
hts01103
B
% of defined area positive for Dlg-Pak PLA signal
22
20
18
16
14
12
10
8
6
4
2
0
*
6
6
wt
hts01103
C
% of defined area positive for Dlg-Drpr PLA signal
16
14
12
10
8
6
4
2
0
***
8
8
wt
hts01103
D
% of defined area positive for Dlg-Drpr PLA signal
12
10
8
6
4
2
0
***
8
8
mef2>wt
mef2>α-specRNAi

FIG S1

FIG S2

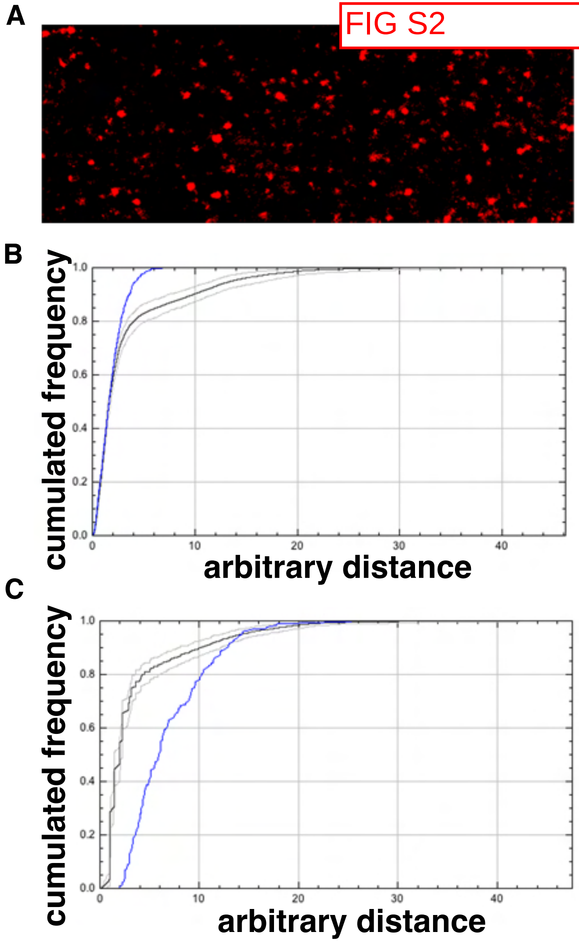
A
B
cumulated frequency
1.0
0.8
0.6
0.4
0.2
0.0
0
10
20
30
40
arbitrary distance
C
cumulated frequency
1.0
0.8
0.6
0.4
0.2
0.0
0
10
20
30
40
arbitrary distance

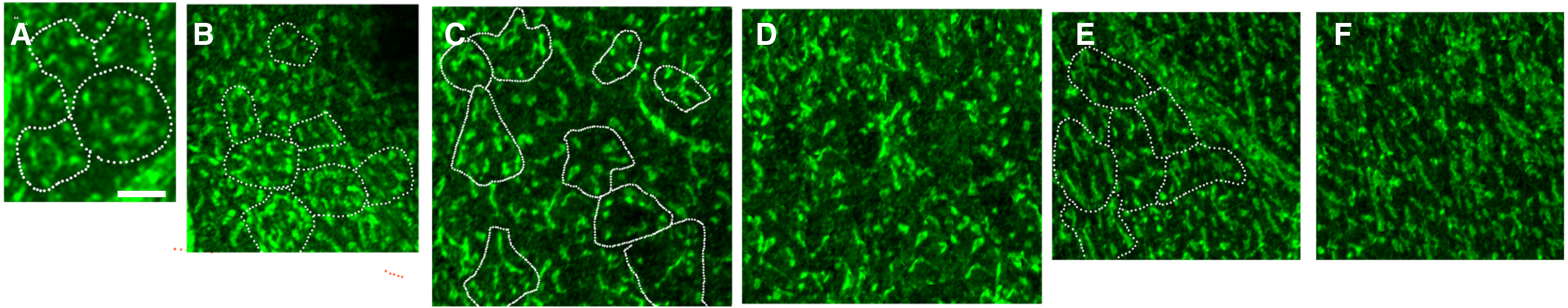


FIG S3

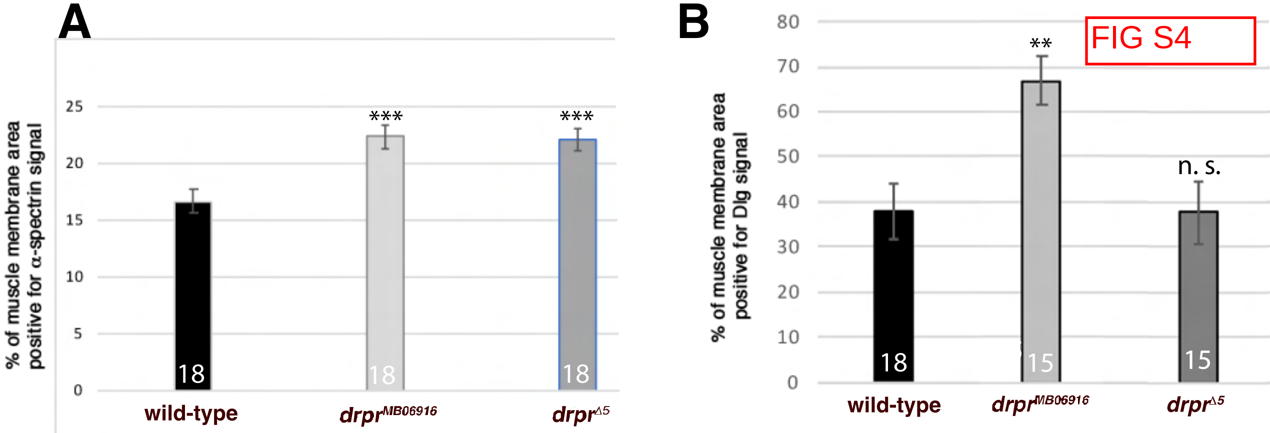

A
% of muscle membrane area positive for α-spectrin signal
0
5
10
15
20
25
***
***
18
18
18
wild-type
drprMB06916
drprΔ5
B
FIG S4
% of muscle membrane area positive for Dlg signal
0
10
20
30
40
50
60
70
80
**
n. s.
18
15
15
wild-type
drprMB06916
drprΔ5

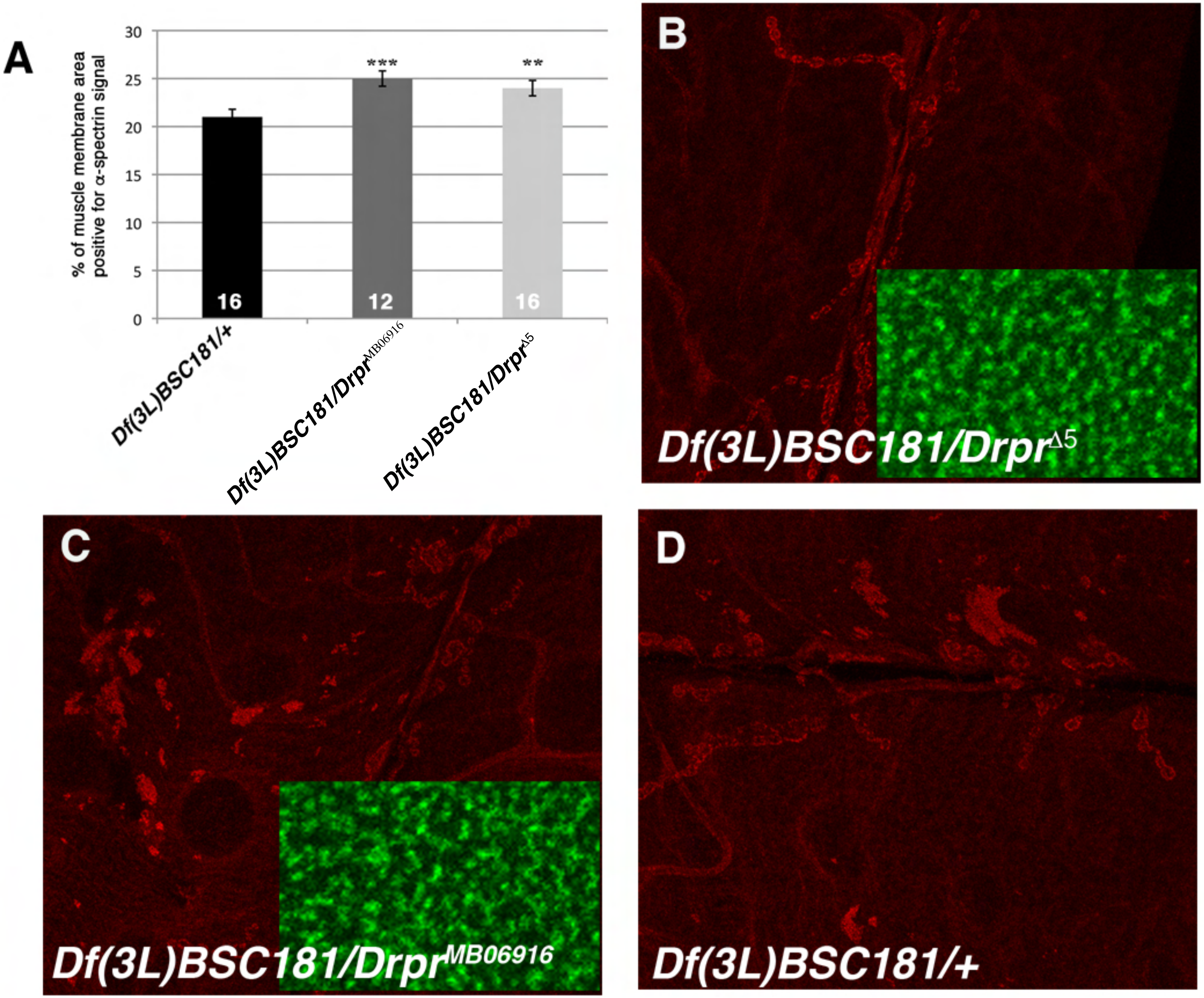
A
% of muscle membrane area positive for α-spectrin signal
30
25
20
15
10
5
0
***
**
16
12
16
Df(3L)BSC181/+
Df(3L)BSC181/DrprMB06916
Df(3L)BSC181/DrprΔ5
B
Df(3L)BSC181/DrprΔ5
C
Df(3L)BSC181/DrprMB06916
D
Df(3L)BSC181/+

FIG S5

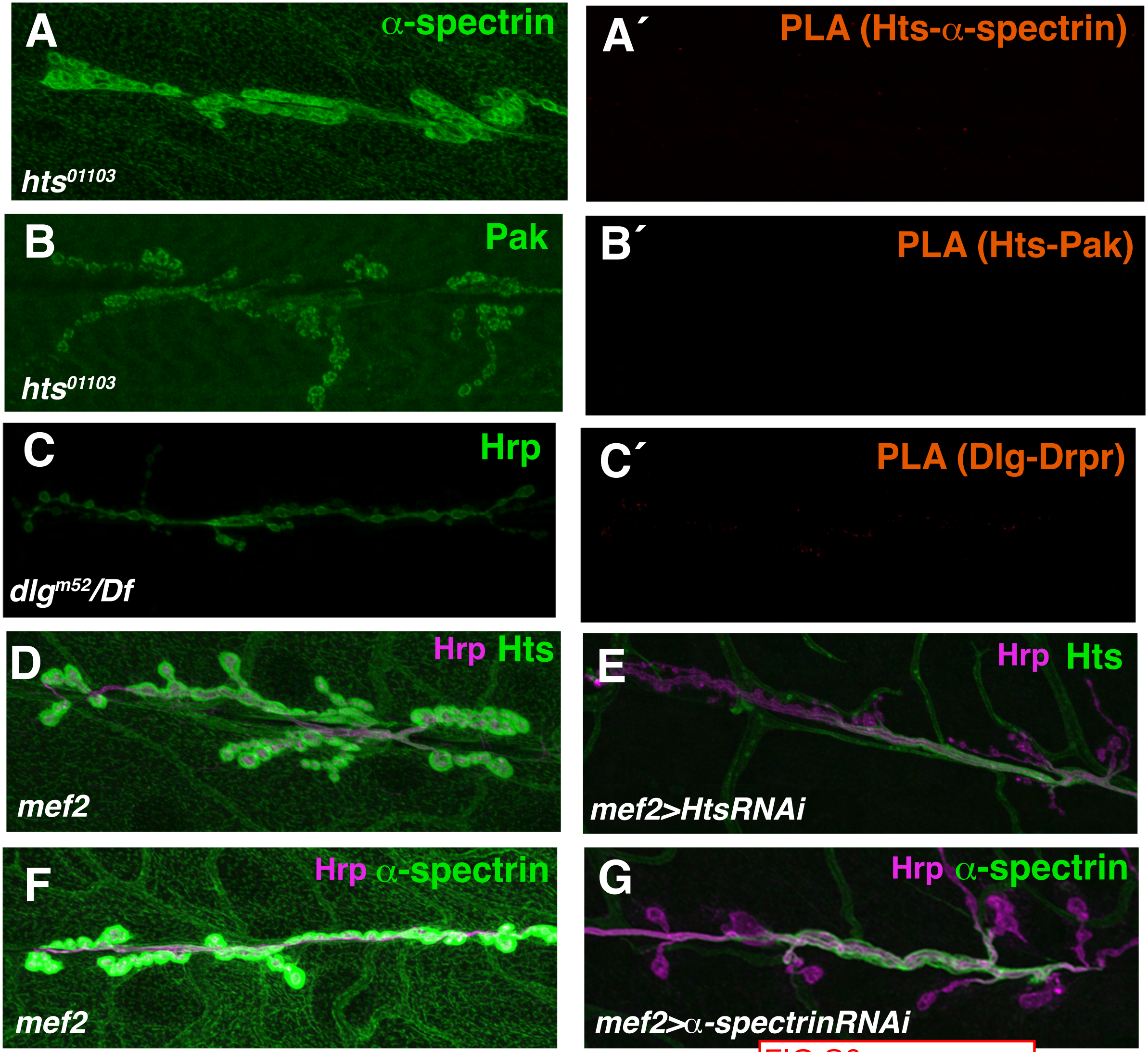
A
α-spectrin
hts01103
A´
PLA (Hts-α-spectrin)
B
Pak
hts01103
B´
PLA (Hts-Pak)
C
Hrp
dlgm52/Df
C´
PLA (Dlg-Drpr)
D
Hrp Hts
mef2
E
Hrp Hts
mef2>HtsRNAi
F
Hrp α-spectrin
mef2
G
Hrp α-spectrin
mef2>α-spectrinRNAi

FIG S6
